# Purkinje cell development and degeneration in the spastic Han-Wistar rat model of ataxia

**DOI:** 10.1101/2025.09.23.677887

**Authors:** Hanna Bellafard, Tanya Pelayo, Michael Aghopoo, Diba Sarhangnejad, Shayn Gaft, Luciana Werneck Zuccherato, Carlos Eduardo G. Amorim, Maria Elena de Bellard

**Affiliations:** Department of Biology, California State University, Northridge, CA, USA

**Keywords:** Purkinje neuron, Motor control, Hereditary ataxia, Rare disease, Whole Genome Sequencing

## Abstract

Hereditary ataxia is a neurodegenerative disorder notable for its early onset, with symptoms appearing in patients as young as two years old. Although affected individuals exhibit severe motor deficits and early mortality rates, the timeline of Purkinje cell loss remains unclear. To address this gap, we used the spastic Han-Wistar rat model, which harbors an unknown homozygous recessive variant that causes Purkinje cell loss. Here, we aimed to determine the onset and temporal progression of Purkinje neuronal loss in the spastic Han-Wistar model. To achieve this, we employed immunohistochemistry, Hematoxylin and Eosin histology, and neuronal density quantification. Behavioral testing demonstrated early-onset, progressive motor impairment in mutant rats, which coincided with a gradual loss of Purkinje cells in the cerebellum. Additionally, guided by pedigree analysis from a previous study indicating autosomal recessive inheritance for this ataxia, we performed whole-genome shotgun sequencing of a parent-offspring trio to identify amino acid-changing mutations consistent with this pattern. We used Sanger sequencing to exclude non-causal candidates. Together, our findings provide new insights into the onset and genetic complexity of ataxia, refining the value of the spastic Han-Wistar rat as a model for investigating mechanisms underlying hereditary ataxia and broader neurodegenerative disorders.

## Introduction

Neurodegenerative diseases represent one of the most pressing challenges in modern medicine, affecting millions of individuals worldwide and imposing substantial burdens on healthcare systems and families^1^. Among these disorders, hereditary ataxias constitute a clinically and genetically heterogeneous group of conditions characterized by progressive cerebellar dysfunction, resulting in impaired coordination, balance, and motor control^2^. These disorders exhibit substantial phenotypic diversity, with onset ranging from early childhood to late adulthood and inheritance patterns spanning autosomal dominant, autosomal recessive, and X-linked transmission^3^.

The human cerebellum, though comprising only 10% of total brain volume, contains more neurons than the rest of the brain combined and serves as a critical hub for motor coordination and cognitive processing^4,5^. Purkinje cells function as the sole output neurons, integrating vast amounts of sensory information through their intricate dendritic arbors and providing inhibitory signals to the deep cerebellar nuclei^6,7^. These neurons are highly specialized yet particularly vulnerable to genetic mutations and environmental insults, with their degeneration serving as a hallmark of cerebellar ataxias^8,9^.

While the selective vulnerability of Purkinje cells in ataxia is well recognized, the temporal dynamics of their degeneration in relation to symptom onset remain unclear, which limits opportunities for early intervention strategies^10^. Furthermore, while numerous genetic loci have been associated with hereditary ataxias, approximately 75% of patients remain without a molecular diagnosis, highlighting the incomplete understanding of the genetic architecture underlying these disorders^11,12^.

Animal models have been invaluable for investigating the mechanisms of cerebellar degeneration and testing potential therapeutic interventions^13,14,15^. The spastic Han-Wistar (sHW) rat represents a proven model of hereditary ataxia, exhibiting progressive motor dysfunction, cerebellar degeneration, and early mortality by approximately 60 days of age (p60)^16^. Previous studies have documented significant Purkinje cell loss and structural abnormalities in the cerebellar cortex; yet, the genetic basis of this mutation remains unknown, and the precise timeline of neurodegeneration has not been systematically characterized^16–18^.

Here, we sought to provide a detailed temporal map of Purkinje cell loss in the sHW rat model, demonstrating that degeneration begins earlier (P15 vs. P21) and precedes the onset of motor symptoms. In addition, we present a comprehensive temporal investigation of Purkinje cell degeneration in the sHW rat model, combining behavioral assessments and histological analyses to address two fundamental questions: (1) What is the temporal progression of Purkinje cell loss relative to the onset of motor symptoms? and (2) What is the progression of motor impairments in mutant rats as measured by rotarod, open-field, and weight analysis? Through Immunohistochemistry (IHC), Hematoxylin and Eosin (H&E) staining, and quantitative neuronal density measurements, we demonstrate that Purkinje cell degeneration begins as early as P15, preceding the emergence of overt motor deficits. Additionally, we employed whole-genome sequencing and Sanger validation to identify candidate genes potentially responsible for the disorder. Our findings provide new insights into the pathological timeline of cerebellar degeneration and contribute to the growing understanding of genetic factors underlying hereditary ataxias.

## Results

This study investigated the progression and timeline of Purkinje cell degeneration in the cerebellum of sHW mutant rats. We employed three experimental approaches: (i) behavioral assays including Open Field, Rotarod, and weight measurement; (ii) histological analyses using H&E staining and IHC; and (iii) whole-genome shotgun sequencing and Sanger sequencing to identify candidate genes underlying the trait.

### Early-onset motor and systemic decline in ataxic rats

Locomotor activity, measured by the open-field test, was significantly reduced in mutant rats compared to wild-type controls. The total distance traveled declined progressively with age (Fig. 1a), reflecting reduced mobility. Wild-type rats traveled an average of 4,190 millimeters compared to 1,846 millimeters in spastic mutants, representing a 56% reduction in locomotor activity (two-way repeated measures ANOVA, p < 0.0001; mean difference = 2,344 units, 95% CI: 1,616-3,072). Motor coordination, assessed using the rotarod test, also declined over time in mutants (Fig. 1b). Wild-type rats remained on the rotarod for an average of 134.6 seconds compared to 73.97 seconds for spastic mutants, representing a 45% reduction in motor coordination performance (two-way repeated measures ANOVA, p < 0.0001; mean difference = 60.65 seconds, 95% CI: 42.24-79.07). Mutant animals exhibited early signs of motor impairment, falling before reaching the traditional target duration of 180 seconds^19^. Deficits became prominent by P35, when most mutants were unable to complete the trial, indicating impaired balance and coordination. Body weight measurements showed a similar progressive decline (Fig. 1c), with significant differences between wild-type and mutant. Weight loss coincided with the onset of motor impairment, suggesting a relationship between systemic health decline and reduced motor performance.

**Figure 1.**
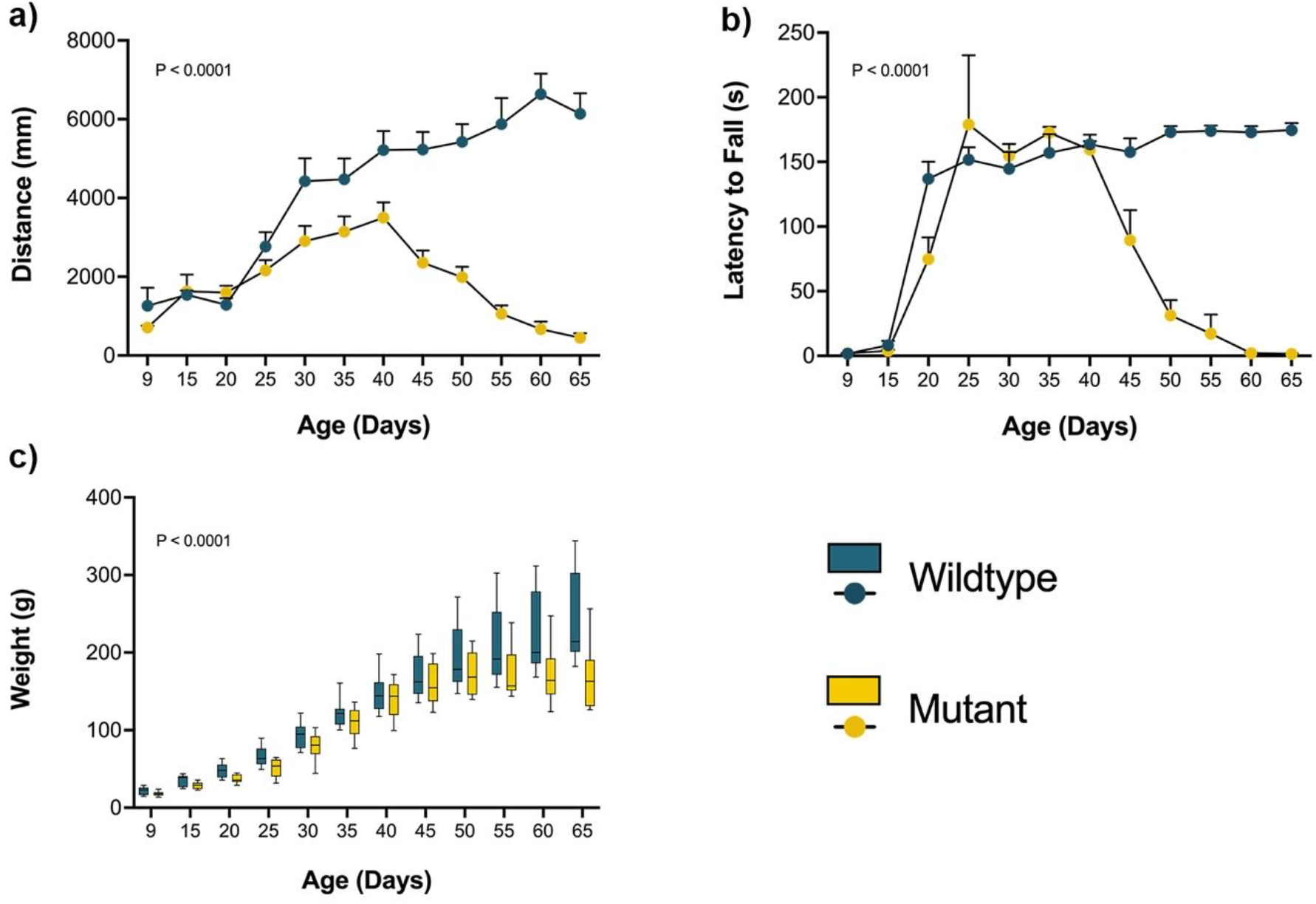
Behavioral assessment of spastic Han-Wistar rats. Behavioral assessments were performed on wild-type and mutant rats (n = 12 per group) every 5 days from P9 to P65. (a) Locomotor activity measured as total distance traveled in the open-field test (120 s duration). (b) Motor coordination assessed by latency to fall in the rotarod test (180 s duration). (c) Body weight measurements over the same period are used to monitor body mass changes. Statistical analysis was performed in all behavioral tests using repeated-measures ANOVA. Error bars represent the standard error of the mean.

### Disrupted morphology and reduced cell counts by H&E

To assess the morphology and physical appearance of Purkinje cells in the cerebellum, we utilized the H&E staining on the cerebellum of sHW mutants and wild-type rats at P15, P30, and P60. Initial observations revealed a noticeable reduction in the number of Purkinje neurons in mutant rats compared to their wild-type counterparts^16^.

In wild-type rats, Purkinje cells exhibited normal morphology, characterized by large, round nuclei (Fig. 2). The cell bodies were neatly aligned in a single layer within the Purkinje cell layer of the cerebellum. The dendritic arborizations were also visible, extending into the molecular layer. In contrast, Purkinje cells in the mutant sHW rats displayed signs of degeneration (Fig. 3a). There was a significant reduction in the number of Purkinje cells, with many neurons appearing shrunken and fragmented (Fig. 2). The alignment of Purkinje cells was disrupted. Quantitative analysis confirmed this reduction, with Purkinje cell loss ranging from 10.4% at P30 to 64.1% at P60, including a 24.6% decrease at P15 (Fig. 3a).

**Figure 2.**
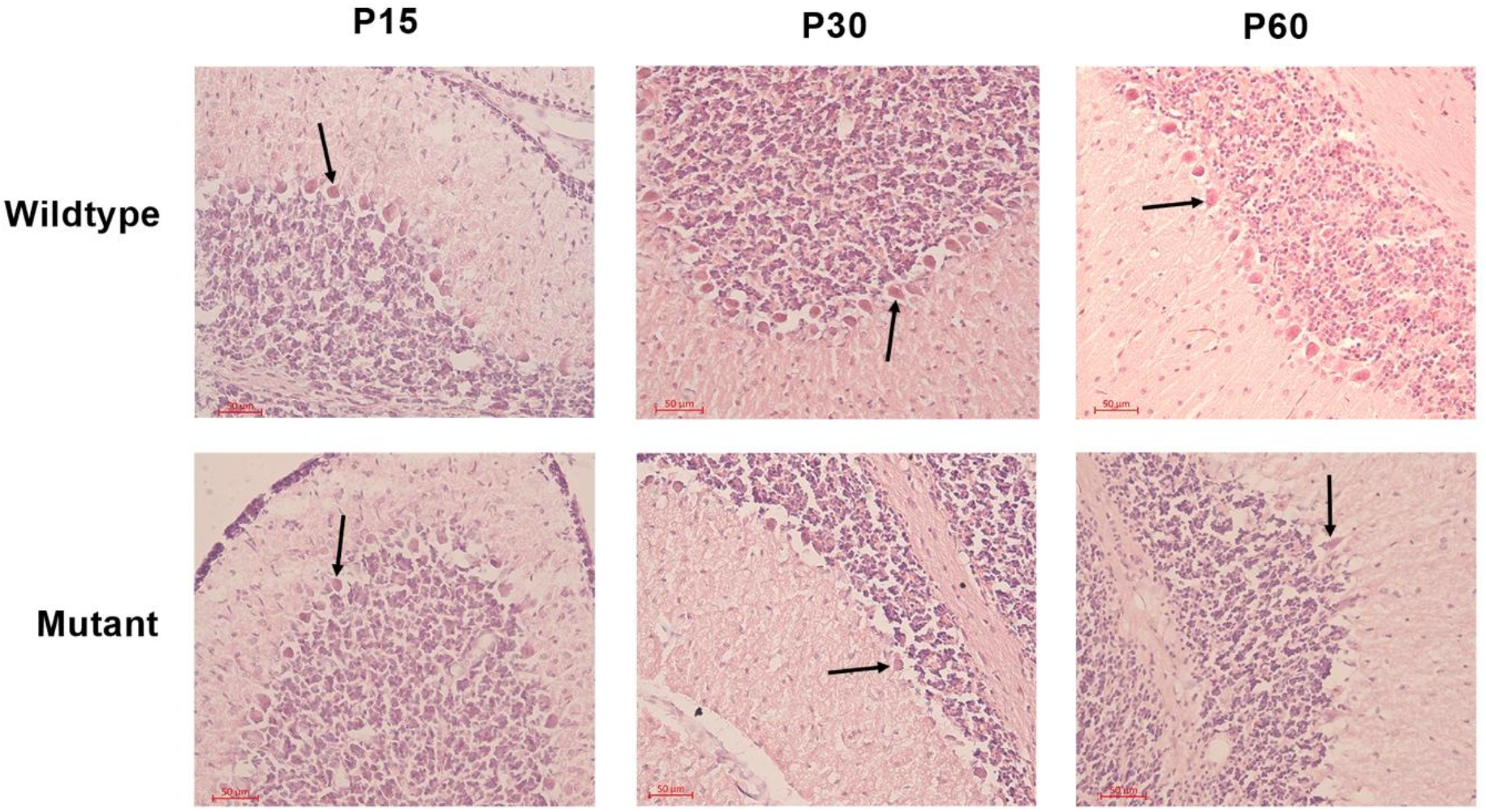
Hematoxylin & Eosin staining of Purkinje cells in the cerebellum of Spastic Han-Wistar rats. Representative H&E-stained sections (50 µm) of wild-type and mutant at P15, P30, and P60. Arrows indicate Purkinje cells in the Purkinje layer.

**Figure 3.**
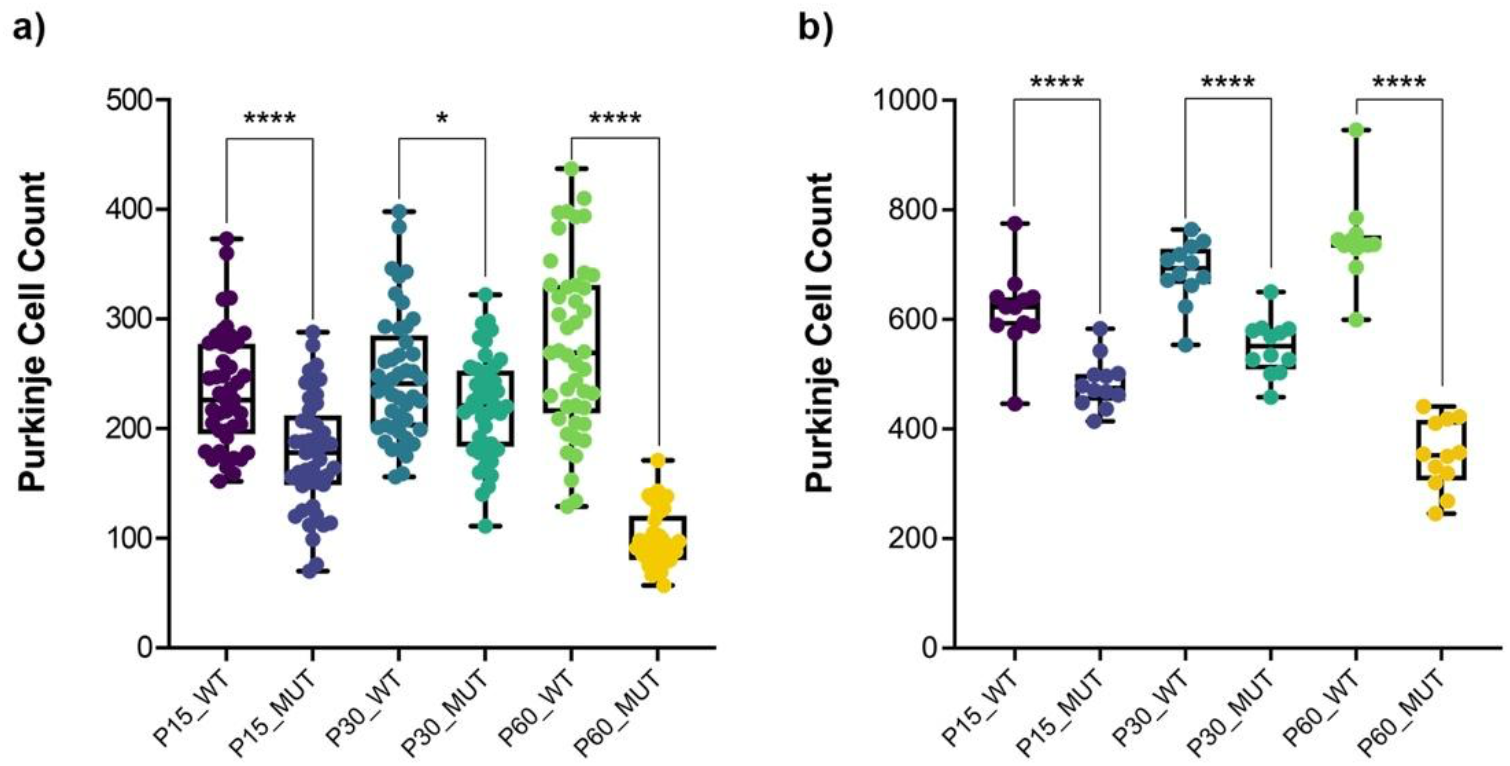
Histological and immunohistochemical quantification of Purkinje cell degeneration. (a) Hematoxylin & Eosin staining quantification of Purkinje cell counts at P15, P30, and P60. Mutant rats showed a significant reduction in Purkinje cells compared to wild-type controls. (b) Immunohistochemical quantification of Purkinje cell counts at P15, P30, and P60. Mutant rats showed a significantly lower number of Purkinje cells compared to wild-type controls. Statistical analysis was performed using Welch’s t-test, with error bars representing the standard error of the mean (SEM).

### Calbindin staining confirms Purkinje cell degeneration

Purkinje cells were visualized using the Calbindin marker in cerebellar sections from wild-type and mutant rats at P15, P30, and P60 (Fig. 4). In wild-type animals, Purkinje cells formed a continuous monolayer with large, well-defined cell bodies and extensive dendritic arborization at all ages examined. Mutant animals had a significantly lower Purkinje cell count than age-matched wild-type controls at all time points, with reductions of 21.5% at P15, 20.0% at P30, and 52.8% at P60. Although the Purkinje cell count in mutants showed an increase from P15 to P30, it remained significantly less compared to wild-type and dropped further by P60 (Fig. 3b). These results parallel the neuronal loss observed in H&E-stained sections and provide cell-type-specific confirmation of Purkinje cell degeneration in the mutant cerebellum.

**Figure 4.**
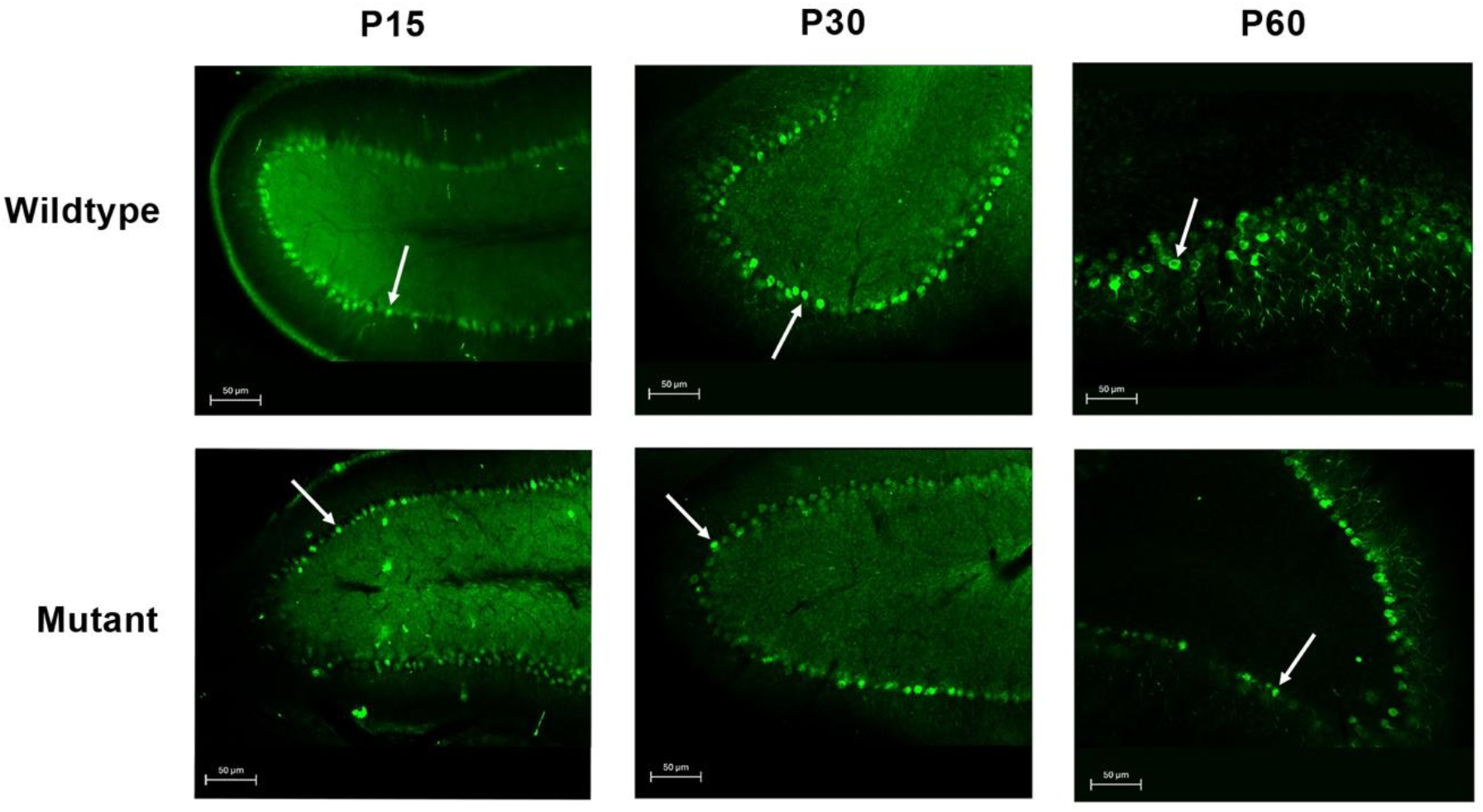
Immunohistochemical analysis of Purkinje cells in Spastic Han-Wistar rats. Representative cerebellar sections (50 µm) from wild-type and mutant rats at P15, P30, and P60. Purkinje cells were labeled with the Calbindin marker. Arrows indicate Purkinje cell bodies.

### Genomic search for candidate genes underlying ataxia in Han-Wistar rats

Guided by previous pedigree analysis indicating autosomal recessive inheritance for this ataxia^17^, we used whole-genome shotgun sequencing (WGS) to search for candidate genes underlying this trait. WGS was performed on a trio from Family 1, consisting of the carrier mother, carrier father, and one affected offspring (Supplementary Fig.1). Genome-wide average depth-of-coverage ranged from ∼23x to ∼25x. Family 1 included three additional siblings, one affected based on phenotypic presentation, and two asymptomatic individuals whose carrier status was unknown at the time of sample collection. These additional siblings were later used for the validation of candidate genes through Sanger sequencing.

Following single-nucleotide variant (SNV) calling, we searched the data for variants following an autosomal recessive inheritance pattern as previously described for this trait^17^. In doing so, we assumed the trait is not caused by a compound heterozygous mutation, which is a common cause of autosomal recessive ataxia^20^. Functional annotation classified the variants into 17 high-impact candidates, 233 moderate-impact, and 24 low-impact, distributed across 273 genes (Supplementary Table 2). To refine this list, candidate prioritization was performed through a systematic review of publicly available resources, including peer-reviewed literature, the NCBI Gene database, the UCSC Genome Browser, and the Human Protein Atlas, to assess gene function, previously reported phenotypes, and tissue-specific expression profiles. This process identified ten genes of interest (Supplementary Table 1), one high-impact gene (*Sbf2*), six moderate-impact genes (*Rhot2, Sh3bp4, Abi2, Cplane1, Stradb, Eif2ak2*), and three low-impact genes (*Pkd1l2, Ranbp9, Caprin2*).

Based on predicted functional impact and biological relevance, the high-impact SNV candidate (chr1:175154510) in the *Sbf2* gene was selected for validation in Family 1 using Sanger sequencing. The segregation pattern was consistent with an autosomal recessive model: both parents were heterozygous for the variant; both phenotypically affected offspring were homozygous for the alternate allele; and the two asymptomatic siblings were confirmed to be one heterozygous carrier and one homozygous for the wildtype reference genotype. To assess whether this variant was conserved across pedigrees, *Sbf2* was subsequently evaluated in an independent family 2 (Supplementary Fig. 1). In this case, genotypes did not segregate with the disease phenotype, and the gene was thus excluded as the causal variant.

In parallel, copy number variant (CNV) analysis was performed on the WGS data. The frataxin (*Fxn*) gene emerged as a notable candidate due to its well-documented role in hereditary ataxia and extensive prior study. *Fxn* was therefore selected for validation in Family 1; however, Sanger sequencing results were inconclusive, and no definitive segregation pattern could be established.

## Discussion

The spastic Han-Wistar rat is a well-established genetic model of hereditary ataxia, caused by a naturally inherited mutation that selectively targets Purkinje cells in the cerebellum. Previous studies suggested that Purkinje cell degeneration emerges around P21, coinciding with the onset of observable motor impairment^17,21^. Our study challenges this assumption by demonstrating that Purkinje cell loss begins earlier, at P15, well before overt behavioral symptoms become apparent. Such early vulnerability is consistent with observations in other ataxic mutants, such as Lurcher and *pcd* mice, where Purkinje cell loss occurs within the first postnatal weeks^22–24^. By refining the temporal onset in sHW rats, our findings address the long-standing question of when cerebellar pathology truly begins relative to symptom onset and provide a more precise disease timeline to guide biomarker discovery and pre-symptomatic intervention strategies.

Behavioral assessments in the sHW rat revealed progressive impairments in locomotion, coordination, and weight gain, with deficits becoming evident around P35. Importantly, these functional changes appeared weeks after histological evidence of Purkinje cell loss, indicating that substantial degeneration can occur before overt symptoms arise. This temporal dissociation has also been documented in spinocerebellar ataxias, where carriers often remain clinically silent despite ongoing neuronal loss^25–27^. A similar threshold-dependent effect is well established in other neurodegenerative disorders, such as Huntington’s disease, where compensatory networks delay clinical onset until a critical proportion of neurons is lost^28^. By extending this principle to the sHW rat, our findings emphasize its utility for modeling the pre-symptomatic phase of hereditary ataxia and for identifying biomarkers that could inform early intervention strategies.

The unresolved genetics of the sHW mutation further underscores the heterogeneity of hereditary ataxias. Our whole-genome sequencing analyses initially highlighted *Sbf2* – the gene associated with Charcot-Marie-Tooth disease in humans – as a strong candidate, with segregation patterns consistent in one pedigree. However, follow-up testing in an independent family did not support this association, leading to its exclusion as a causal variant. Similarly, copy number variation analysis suggested a possible role for *Fxn*, a gene implicated in Friedreich’s ataxia in humans, but validation results were inconclusive. Notably, recent discoveries such as the identification of pathogenic GAA repeat expansions in the *FGF14* gene, associated with SCA27B ataxia in humans, demonstrate that Next-Generation sequencing approaches, based on short sequencing reads, may fail to detect repeat expansion disorders, underscoring an additional challenge in resolving undiagnosed cases^29^. Together, these findings highlight the genetic complexity of hereditary ataxia and support the clinical observation that nearly 75% of patients remain without a molecular diagnosis, despite advances in sequencing^11,12,30^. Rather than representing a limitation, this process of iterative exclusion narrows the pool of candidates and mirrors the diagnostic pathway in human patients, where excluding false leads is often as critical as identifying the causative mutation^31^. By refining the genetic search space, our work provides a valuable resource for future studies with larger cohorts and more comprehensive sequencing approaches to pinpoint the underlying variant.

The identification of early Purkinje cell degeneration in the sHW rat also has important implications beyond genetics. Establishing a cellular timeline that precedes behavioral onset provides a foundation for identifying early biomarkers of disease progression. Such markers are crucial for designing interventions during the pre-symptomatic stage, when treatments are most likely to be effective. Moreover, clarifying the onset of degeneration aligns the sHW rat more closely with human ataxias, where cerebellar atrophy is detectable years before motor symptoms appear^25,27^.

Several limitations of this study should be acknowledged. The genetic analyses were conducted on small pedigrees, limiting the power to confirm segregation across multiple families. Larger, multi-generational cohorts will be essential to establish inheritance patterns more conclusively. In addition, although our histological data reveal degeneration by P15, finer sampling across earlier developmental ages will be necessary to determine whether Purkinje cell loss begins even sooner.

## Conclusion

In conclusion, our study demonstrates that Purkinje cell degeneration in the sHW rat begins earlier than previously recognized, preceding the onset of motor deficits by several weeks. Behavioral decline closely follows cellular loss, reinforcing the central role of Purkinje neurons in ataxic pathology. In parallel, our genetic analyses excluded two high-priority candidate genes. We also present a curated list of 273 genes harboring mutations consistent with autosomal recessive inheritance, offering a valuable resource for ongoing efforts to identify the causal variant. Together, these findings establish a detailed temporal framework for cerebellar degeneration in this model and highlight its utility for investigating the early stages of hereditary ataxia and for testing pre-symptomatic therapeutic strategies.

## Methods

### Animals

Spastic Han-Wistar rats were obtained from the California State University, Northridge (CSUN) Vivarium breeding colony between the years 2023 and 2024. Both male and female rats were used, as no sex differences in phenotype have been reported in this model^17^. Animals were housed in groups of 2–4 and provided unrestricted access to standard rat chow and water. Housing was maintained on a 12 h light–dark cycle at 23–25 °C. All procedures were conducted in accordance with CSUN’s Institutional Animal Care and Use Committee (IACUC) guidelines.

### Breeding mutants

Upon weaning, animals were screened for symptoms of the mutant phenotype at approximately p15. Mutants were confirmed based on observable traits such as hyperactivity and severe whole-body tremors. Additionally, mutants tended to exhibit decreased body weight and weaker grip strength compared to their wild-type siblings.

### Behavioral analysis

The behavioral component of this study aimed to evaluate motor coordination and activity levels in 12 mutant and 12 wild-type rats at various developmental stages. Animals aged 9 days old were selected for assessment. Firstly, they underwent the rotarod test. Each trial was conducted at 10 RPM for a maximum of 180 seconds and repeated three times per subject. Subsequently, the rats were placed in a 100 × 100 cm box made of black acrylonitrile butadiene styrene (ABS) plastic. This test consisted of placing the rodent in the described box three times, each time for 120 seconds, during which they were free to move about and explore their environment. The animals were given a 120-second break between each run. All actions within this chamber were captured with a Logitech Quick-Cam Pro 90000 (ffdshow codec 1:122 compression ratio, 15 fps) and recorded to a computer hard drive using VirtualDub. Video tracking and movement analysis were conducted using custom scripts written in Python with the OpenCV computer vision library (opencv.willowgarage.com). Lastly, the weight of each animal was recorded to monitor potential correlations between behavioral performance and physical development. These three tests were conducted every 5 days, starting when the rats were 9 days old and continuing until they reached 65 days of age.

### Brain fixation and harvesting

At the designated age, the animals were euthanized, and the brains were extracted for analysis. This process involved exposing the animals to CO_2_ followed by a transcardiac perfusion. Animals were placed in a perfusing pan, and a midline incision through the sternum was made to expose the heart for upper body perfusion using transcardiac puncture. Using hemostats, the descending aorta was clamped to prevent whole-body perfusion. A 26-gauge needle was inserted into the left ventricle, the right atrium was then immediately incised, and a peristaltic pump was activated with a flow rate of 7 ml per minute. Animals were perfused with a 0.9% saline solution to clear most of the blood from the head and neck region. The paleness of the eyes, ears, and forelimbs visually confirmed clearance of blood, and the stiffness of the neck further confirmed clearance of blood. Afterward, 4% paraformaldehyde at pH 7.4 was used on the animal tissue until complete fixation was achieved. The perfusion is also confirmed by physical inspection of increased stiffness in the neck. The brain was then removed, placed in 4% paraformaldehyde, and stored at 4 °C for future use.

### Tissue sectioning and mounting by cryostat

Cerebellums of harvested brains were sectioned using a cryostat microtome, Leica CM1950, and mounted on a cryostat-mounting platform using tissue freezing media (Tissue-Tek OCT compound 4583). These brains were then sectioned at −20 °C into 15μm sagittal cuts. After sectioning, the sections were promptly mounted onto superfrost slides and allowed to dry for 24-48 hours in a 4°C refrigerator to ensure proper adhesion before staining. For each animal, fifteen randomly selected tissue sections were used for H&E staining.

### Hematoxylin and eosin staining

The morphology and structural integrity of Purkinje cells were evaluated using Harris’s hematoxylin and eosin (H&E) staining. Cerebellar sections from mutant rats and wild-type littermates were processed in parallel. Sections were rehydrated through graded ethanol (100% and 95%), immersed in hematoxylin for five dips (with staining duration adjusted to achieve optimal intensity), and rinsed in tap water. Slides were differentiated in 1% acid alcohol (1 mL HCl in 99 mL 95% ethanol), rinsed in 2% aqueous sodium bicarbonate to enhance nuclear staining, and counterstained with eosin Y for 20 s. Following dehydration in graded ethanol, slides were cleared in xylene and coverslipped using Permount. Purkinje cell morphology and evidence of neuronal degeneration were assessed in mutants at postnatal days 15, 30, and 60 using light microscopy.

### Tissue embedding and vibratome sectioning

For IHC, cerebella were embedded in 4% agarose prepared in 1× phosphate-buffered saline (PBS). To prepare the agarose, 4 g of agarose powder was added to 100 mL 1× PBS, allowed to hydrate at room temperature, and then heated in a microwave oven for ∼2 minutes with intermittent swirling every 30 s to prevent boiling and foaming, until fully dissolved. The solution was cooled to approximately 40 °C before embedding to avoid heat damage to the tissue. Tissues were placed in appropriately sized disposable plastic molds, and molten agarose was gently poured to fully submerge the tissue, ensuring it was lifted slightly so that a thin agarose layer separated the tissue from the mold base. Once the agarose had solidified at room temperature, blocks were wrapped in a damp paper towel, sealed in plastic bags, and stored at 4 °C for up to 24 hours before sectioning.

For vibratome sectioning, excess agarose surrounding the tissue was trimmed, and blocks were mounted with the desired orientation maintained. Sagittal sections of cerebellum were cut at 100 μm thickness using a vibratome (Leica, VT1000S) according to the manufacturer’s instructions. Sections for IHC were collected into wells containing 4% paraformaldehyde (PFA) for storage before staining.

### Immunohistochemistry

Free-floating cerebellar sections were processed for IHC staining to visualize Purkinje neurons. Sections were transferred into new wells and blocked with 10% goat serum in phosphate-buffered saline (PBS) for 4 hours at 37 °C to prevent nonspecific binding. Following blocking, the tissues were rinsed twice with 1× PBS and then incubated in 600 µL PBS containing the anti-Calbindin primary antibody (4 µL per well), a marker for Purkinje cells (*1:500, Abcam, Cat# ab108404)*. Sections were incubated for 48 h at 37 °C. After primary antibody incubation, sections were washed six times with PBS (30 min each) on a gentle shaker, adding PBS along the side of the well to minimize tissue disruption. Samples were then incubated overnight at 37 °C with Alexa Fluor 488 conjugated anti-rabbit (4 µL/well) secondary antibodies. The following day, sections were washed six times with PBS (30 min each) using the same gentle washing method. Sections were mounted on slides, coverslipped, and imaged using a Zeiss fluorescence microscope equipped with appropriate filter sets for Alexa Fluor 488 (*1:500, Invitrogen, Cat# A-11008)*.

Images were acquired at consistent exposure times, saved as high-resolution digital files, and processed using AxioImage software (Zeiss).

### Genetic mapping

A family trio was selected for whole-genome sequencing (WGS), comprising carrier parents, approximately one year of age, and an affected offspring. Both parents were heterozygous carriers of the spastic Han-Wistar allele, as they had previously produced progeny with and without the disease phenotype^17^. In the litter from which the trio was derived, an additional two affected and two unaffected offspring were identified at 60 days of age for further DNA extraction and Sanger confirmation of potential candidate genes. Pedigrees are shown in Supplementary Figure 1. Total DNA was extracted from liver tissue samples obtained from mutants and wild-type rats using the DNeasy Blood & Tissue Kit (Qiagen, Germany), following the manufacturer’s protocol.

Raw sequencing reads were processed using the Illumina DRAGEN Germline Pipeline (v4.3.13), aligned to the *Rattus norvegicus* reference genome (Rnor_6.0). The DRAGEN pipeline performs key steps, including read mapping, sorting, duplicate marking, and germline variant calling. In each step, the default parameters from the DRAGEN pipeline were used. Quality control metrics were summarized using MultiQC. Variant Call Format (VCF) files generated from Germline analyses were annotated using the Ensembl Variant Effect Predictor (VEP, v113) with the Ensembl RN6.104 annotation set. VEP was used to determine the predicted functional consequences of each variant. CNV calling was performed using the previously generated BAM files and the DRAGEN Germline pipeline (v4.3.13). Segmentation was conducted using the Shifting Level Model.

Candidate gene analysis was performed by filtering both annotated SNV and CNV datasets to identify variants following an autosomal recessive inheritance pattern. For SNVs, variants where the father and mother exhibited heterozygous genotypes (0/1) while the proband displayed homozygous alternate genotypes (1/1) were extracted. For CNVs, only deletions following the same genotype pattern were considered, with additional copy number requirements of copy number = 1 in both parents and copy number = 0 in the proband. Functional impact prioritization was performed using VEP impact predictions, with variants classified as HIGH impact (transcript ablation, stop gained, stop loss, frameshift variants) and MODERATE impact (missense variants, in-frame indels) prioritized for downstream analysis. The resulting candidate gene list represented loci harboring putative pathogenic recessive variants that could explain the observed ataxia phenotype in the proband. All filtering and analysis steps were performed using bedtools^32^ and custom shell scripts.

### Candidate gene validation

Primers were designed for Polymerase Chain Reaction (PCR) experiments based on candidate genes identified from the WGS results using Primer3 software (https://primer3.ut.ee/). PCR reactions were performed using 100 ng of genomic DNA in a reaction mixture containing 1× buffer, 4 mM MgCl_2_, 0.2 mM dNTPs, 0.4 μM forward and reverse primers, and 0.05 U/μL Taq DNA polymerase. Thermal cycling conditions were optimized as follows: an initial denaturation at 95 °C for 2 min; 30 cycles of denaturation at 95 °C for 30 s, annealing at 60 °C for 30 s, and extension at 72 °C for 1 min; and a final extension at 72 °C for 7 min. The primers used were as follows: *Sbf2* forward: 5′-ACATCTCAGCACCCGCTTCC-3′, reverse: 5′-TCACAGTAGCACGGAATTTGA-3′; *Fxn* forward: 5′-ATCTGATGCCCCTCTTCTGG-3′, reverse: 5′-TGGGAGTGAATTCAGGGCTT-3’.

PCR products were purified using the ExoSAP-IT Express kit (Thermo Scientific, USA) according to the manufacturer’s protocol. Sequencing reactions were prepared in a final volume of 10 μL using the BigDye Terminator v3.1 Cycle Sequencing Kit (Thermo Scientific, USA), with 1 μL of purified PCR product and 1 μM of either forward or reverse primer. Thermal cycling for sequencing consisted of an initial denaturation at 96 °C for 1 min, followed by 25 cycles of 96 °C for 10 s, 50 °C for 5 s, and 60 °C for 45 s, with a ramp rate of 1 °C/s. Sequencing products were purified using the BigDye XTerminator Purification Kit (Thermo Scientific, USA) and analyzed on a SeqStudio Genetic Analyzer (Thermo Scientific, USA). The resulting sequences were aligned to the *Rattus norvegicus* reference genome assembly Rnor_6.0 (rn6) using Geneious R9 software, and genotypes for the selected genes were annotated accordingly.

## Data availability

Raw whole-genome sequencing data (FASTQ files) from the parent-offspring trio have been deposited in the NCBI Sequence Read Archive (SRA) under BioProject accession number *[to be provided upon acceptance/submission]*. The other datasets generated during the current study are available from the corresponding author upon reasonable request.

## Acknowledgements

We thank George Radionov, Shanna Aghasi, Melesys Kozian, Chanel Jacobo, Michelle Bonilla, and Toni Uhlendorf for their assistance with behavioral testing and histology. We are also grateful to Dr. Randy Cohen for his thoughtful review of the manuscript and to Dr. Cheryl Van Buskirk for her valuable input on the research design during the early stages of this project. We thank the CSUN vivarium members, Jackie Lopez and Elizabeth Perez, for providing maintenance of rats. Finally, we thank members of the De Bellard Lab and Amorim Lab for their support and helpful discussions throughout the course of this project.

## Authors contributions

HB performed the behavioral, histological, and molecular biology experiments and wrote the manuscript with input from all authors. MA, DS, and SG contributed to histological sample preparation and imaging. TP and LWZ conducted the genomic analyses. HB and TP prepared the figures and visual representations of the data. CEGA supervised the genomic component of the study. MEdB conceptualized and supervised the overall project. All authors reviewed and approved the final version of the manuscript.

## Ethical approval

All animal procedures were approved by the Institutional Animal Care and Use Committee (IACUC) at CSUN (protocol #2427-001) and were conducted in strict accordance with the guidelines of the [Include the specific national or international guidelines followed, e.g., “NIH Guide for the Care and Use of Laboratory Animals”.

## Supplementary Information

**Supplementary Figure 1**. Pedigrees for candidate gene validation. In Family 1, whole-genome sequencing was performed on the mother, father, and one affected offspring. Sanger sequencing of the *Sbf2* gene was performed for all individuals in both families.

**Supplementary Table 1**. Candidate genes identified from WGS and filtered by predicted impact and disease relevance.

**Supplementary Table 2**. Functional annotation classified the variants into 273 genes (high, moderate, low).

