## Supplementary figures and images for "Purkinje cell development and degeneration in the spastic Han-Wistar rat model of ataxia"

### Supplementary Figure 1

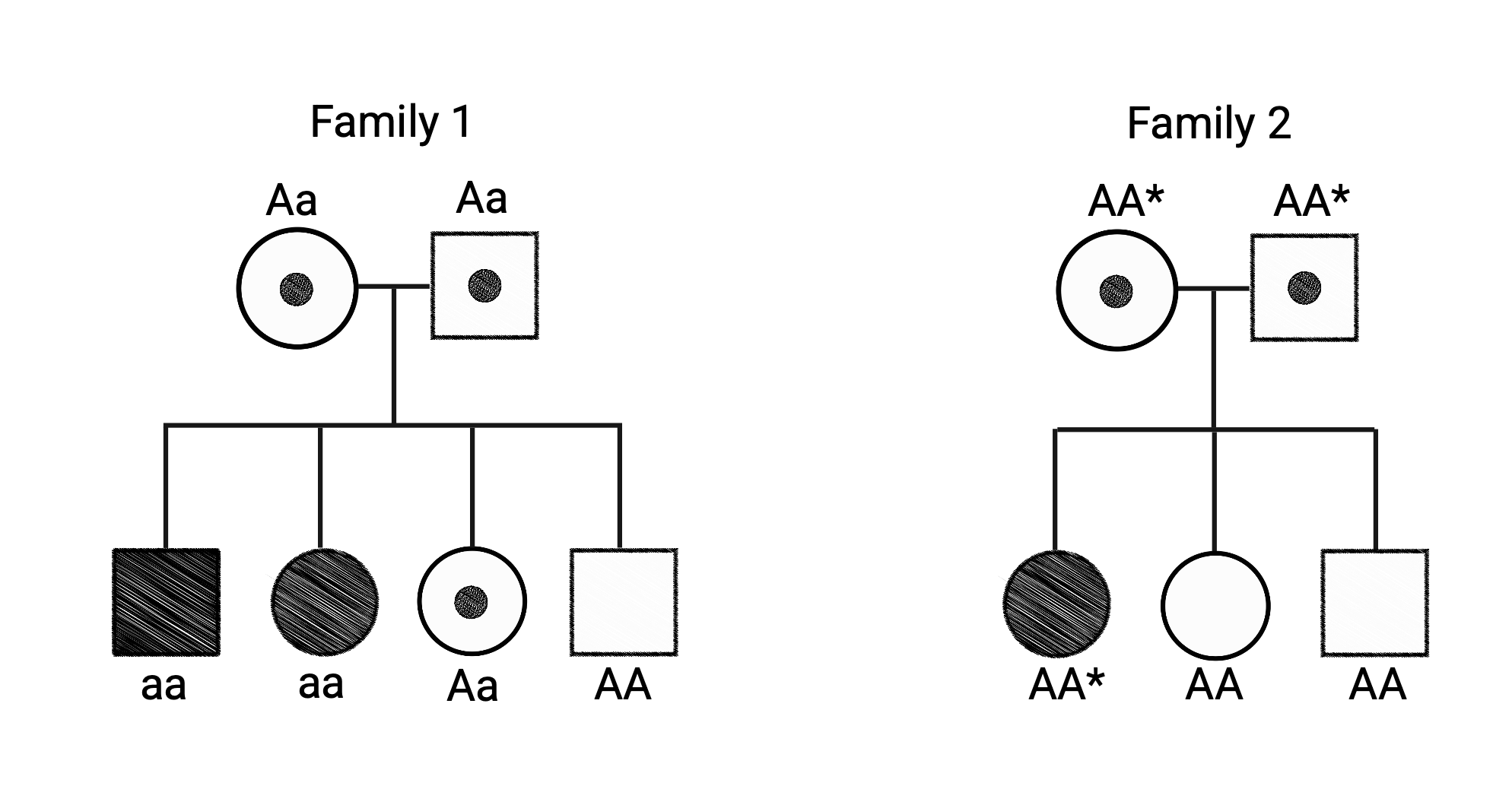
